# Physiological and life history changes associated with seasonal adaptation in the cactophilic *Drosophila mojavensis*

**DOI:** 10.1101/2021.11.09.447153

**Authors:** Tierney M. Shaible, Luciano M. Matzkin

## Abstract

Many insects inhabiting temperate climates are faced with changing environmental conditions throughout the year. Depending on the species, these environmental fluctuations can be experienced within a single generation or across multiple generations. Strategies for dealing with these seasonal changes vary across populations. *Drosophila mojavensis* is a cactophilic *Drosophila* species endemic to the Sonoran Desert. The Sonoran Desert regularly reaches temperatures of 50°C in the summer months. As individuals of this population are rare to collect in the summer months, we therefore simulated in a temperature and light controlled chamber the cycling temperatures experienced by *D. mojavensis* in the Sonoran Desert from April to July (four generations) to understand the physiological and life history changes that allow this population to withstand these conditions. In contrast to our hypothesis of a summer aestivation, we found that *D. mojavensis* continue to reproduce during the summer months, albeit with lower viability, but the longevity of the population is highly reduced during this period. As expected, stress resistance increased during the summer months in both the adult and the larval stages. This study examines several strategies for withstanding the Sonoran Desert summer conditions which may be informative in the study of other desert endemic species.

## 1. Introduction

The majority of organisms are subject to seasonally changing factors over the course of the year, especially those outside of the tropical belt (Roberts, 1978). In many cases, seasonal changes lead to an unfavorable period for survival and/or reproduction. Changes in temperature, humidity, day length, resource availability, predator abundance, etc., are all likely to occur as seasons change throughout the year (Masaki, 1980; Saulich and Musolin, 2017). Changes to these variables are expected to elicit a multifaceted phenotypic response. The response to these climatic fluctuations can be enacted through several different mechanisms, the cost of which, in most cases, is not well understood (Snell-Rood et al., 2018). These environmental changes require organisms to generally respond with one of three main strategies: organism can migrate to more favorable locations, express a seasonally adapted plastic response, or enter a period of dormancy (Roberts, 1978; Tauber et al., 1985; Huestis and Lehmann, 2014).

In desert environments, such as in the Sonoran Desert, abiotic conditions can vary drastically, with temperature ranging from 5-50°C and relative humidity fluctuations from 8% - 80% (Gibbs et al., 2003; Contreras et al., 2013; see Results). Among the many resident species exposed to these extreme and variable environmental conditions is the cactophilic fly, *Drosophila mojavensis*. This species is composed of four ecologically and genetically distinct populations inhabiting the desert regions of the southwest United States and northwest Mexico (Patterson and Crow, 1940; Heed, 1978; Ruiz et al. 1990). *Drosophila mojavensis* utilize necrotic cactus tissues to oviposit therein, develop, and feed as both larvae and adults (Heed, 1978). These flies are largely feeding on the yeast and bacteria that grow within the rots (Ruiz and Heed, 1988). The necrotic cactus host of *D. mojavensis*, with its distinct set of both toxic and nutritional compounds, in addition to abiotic factors (humidity and temperature), has shaped the evolutionary trajectory of this species (Kircher and Bird, 1982; Fogleman and Danielson, 2001; Matzkin, 2014; Allan and Matzkin, 2019). In the Sonoran Desert, *D. mojavensis* utilizes necrotic organ pipe (*Stenocereus thurberi*) cactus as its main host (Heed 1978; Ruiz et al. 1990).

During the thermally stressful summer months, collection of *D. mojavensis* from the Sonoran Desert is rare (Breitmeyer and Markow, 1998; T.M. Shaible pers. obs.). Migration away from their home range is unlikely, as *D. mojavensis* depends on their necrotic cactus host for survival and reproduction (Ruiz and Heed, 1988), and *S. thurberi* is not found outside of the range of the stressful heat and humidity of the Sonoran Desert (Yetman, 2006). Additionally, numerous studies have reported little genetic flow between the populations of *D. mojavensis* (Matzkin, 2004; Reed et al., 2007; Machado et al., 2007; Matzkin, 2008; Pfeiler et al., 2009; Smith et al, 2012).

The environmental conditions experienced by Sonoran Desert endemics have shaped their evolution and can be observed by the specializations these species have. *Drosophila mojavensis* for example, exhibits high levels of thermal and desiccation resistance compared to more mesic adapted *Drosophila* (Stratman and Markow 1998; Krebs, 1999, Gibbs and Matzkin, 2001; Gibbs et al., 2003). Given that migration out of the Sonora Desert does not likely occur in *D. mojavensis*, these types of plastic physiological responses could explain how the species endures the Sonoran Desert summers. One possibility is that the species could be displaying a seasonally adapted plastic response or state of dormancy, such as aestivation, that is the result of the changing environmental conditions approaching the summer months.

Aestivation is characterized by induced suites of preprogrammed changes in the organism due to environmental factors such as day length and temperature (Masaki, 1980; Saulich and Musolin, 2017). The process can include changes to metabolism, reproduction, tolerance to stressors, longevity, and gene expression (Masaki, 1980; Saulich and Musolin, 2017). Aestivation is widespread phenomena in insects and has been observed in 183 cases across 13 orders (Masaki, 1980). While aestivation has not been shown in *Drosophila* species, diapause has been observed in *D. melanogaster* (Schmidt et al., 2005). Although there is speculation on the mechanistic differences between diapause and aestivation, they are typically thought to be under the control of similar sets of pathways (Huestis and Lehmann 2014; Koštál et al. 2017). As there are similar changes and outcomes between the two, these shared pathways are likely conserved across processes, especially considering that some species carry out both processes at different times (Huestis and Lehmann, 2014). Among others life history and physiological effects, an often consequence of the expression of the diapausing phenotype in insects is the extension of life span (Huestis and Lehmann, 2014). Hence, if aestivation is the mechanism used in *D. mojavensis*, it would be expected that it would reallocate resources out of reproduction, restrict activity level, and reduce metabolic activity to increase stress resistance, increase nutritional reserves, and increase longevity. Under such scenario Sonoran *D. mojavensis* would survive the extreme desert environment as adults and resume reproduction once temperature decreases in the fall. The mean adult lifespan for this species is reported to be approximately three weeks (Jaureguy and Etges, 2007), although in benign lab conditions the maximum lifespan can be much longer, > 50 days (Etges and Heed, 1992; T. M. Shaible, pers. obs.). Regardless, an individual’s longevity would need to be significantly extended for an individual to persist past the thermal peak of the summer months.

Without the possibility to easily observe these flies in the field in the summer, we chose to take a multi-generational approach in which we simulated the Sonoran Desert spring and summer field conditions, from April to July. By doing so, we wanted to capture plastic and transgenerational effects on behavioral and physiological traits that happen in response to both simulated field temperatures and photoperiod in the lab. Each generation, we tested phenotypic traits including reproductive output, longevity, various stress responses, activity levels, and nutritional makeup of individuals. By measuring these phenotypic traits, we aimed to test whether *D. mojavensis* that were raised in summer conditions would have an alternative plastic response to those conditions, or instead would cease reproductive output and increase longevity, indicating a canonical aestivation response.

## 2. Methods

### 2.1 Cactus necrosis field temperature

From May 15^th^ to October 19^th^, 2007 a total of 11 temperature data loggers (Onset) were attached to organ pipe cacti at Organ Pipe National Monument in the Sonoran Desert. Three of these data loggers had additional probes which were inserted into necrotic organ pipe cactus.

Internal necrotic cactus and external ambient temperatures were recorded every hour during this period (∼3760 temperature measurements per logger).

### 2.2 Drosophila population

All *D. mojavensis* used in this study were from a multifemale outbred population. For the mass populations 20 isofemale lines, collected from Organ Pipe National Monument in February 2018 were pooled into a population cage (30cm x 30cm x 30cm; BugDorm) and allowed to interbreed for four generations. An outbred population was maintained in a Percival incubator in a 14:10 light:dark cycle at 25°C and 50% relative humidity. From the mass population, eggs were collected for subsequent experiments. A petri dish containing banana-molasses food (Coleman et al., 2018) topped with baker’s yeast to promote egg laying was placed in the cage for 24 hours, after which eggs were transferred in groups of 40 into 30 mL vials containing banana-molasses food.

### 2.3 Experimental setup

Eggs were collected for the first experimental generation in banana-molasses food plates as described above over three days, with a new plate each day. From these plates, 40 vials of about 40 eggs/vial were collected and put into the first experimental treatment temperature cycle (April); another 40 vials of about 40 eggs/vial were collected over the same three days and kept in a 25°C temperature chamber. The experimental treatment temperature cycles are an hourly temperature cycle based on average monthly temperatures for each month from April through July over the years 2011 – 2013 (NCEI: NOAA) (see Fig. 1). The experimental treatment daylight cycles were matched to the mid-month day-length times of 2012 for each month, taken from the Organ Pipe National Monument Visitor’s Center (“Sunrise Sunset Calendars”) (Fig. 1). The daylight cycle for the 25°C temperature chamber was maintained at a constant 14:10 light:dark cycle. Eggs collected for the experimental temperature cycle became the experimental temperature population (ETP), while the eggs collected for the 25°C temperature chamber became the constant 25°C population (CON) (see Fig. 2). Eclosed flies from these vials were collected over a five-day period and transferred into a cage. They were maintained in the cage at their respective temperature cycle for 10 days after the last adult collection occurred. On the 10^th^ day, eggs were collected from a plate within each cage (as described above). The eggs were transferred into the next generation’s temperature cycle. After egg collection, the adult flies within the cage were used for the life history, behavioral and metabolic assays.

**Fig 1.**
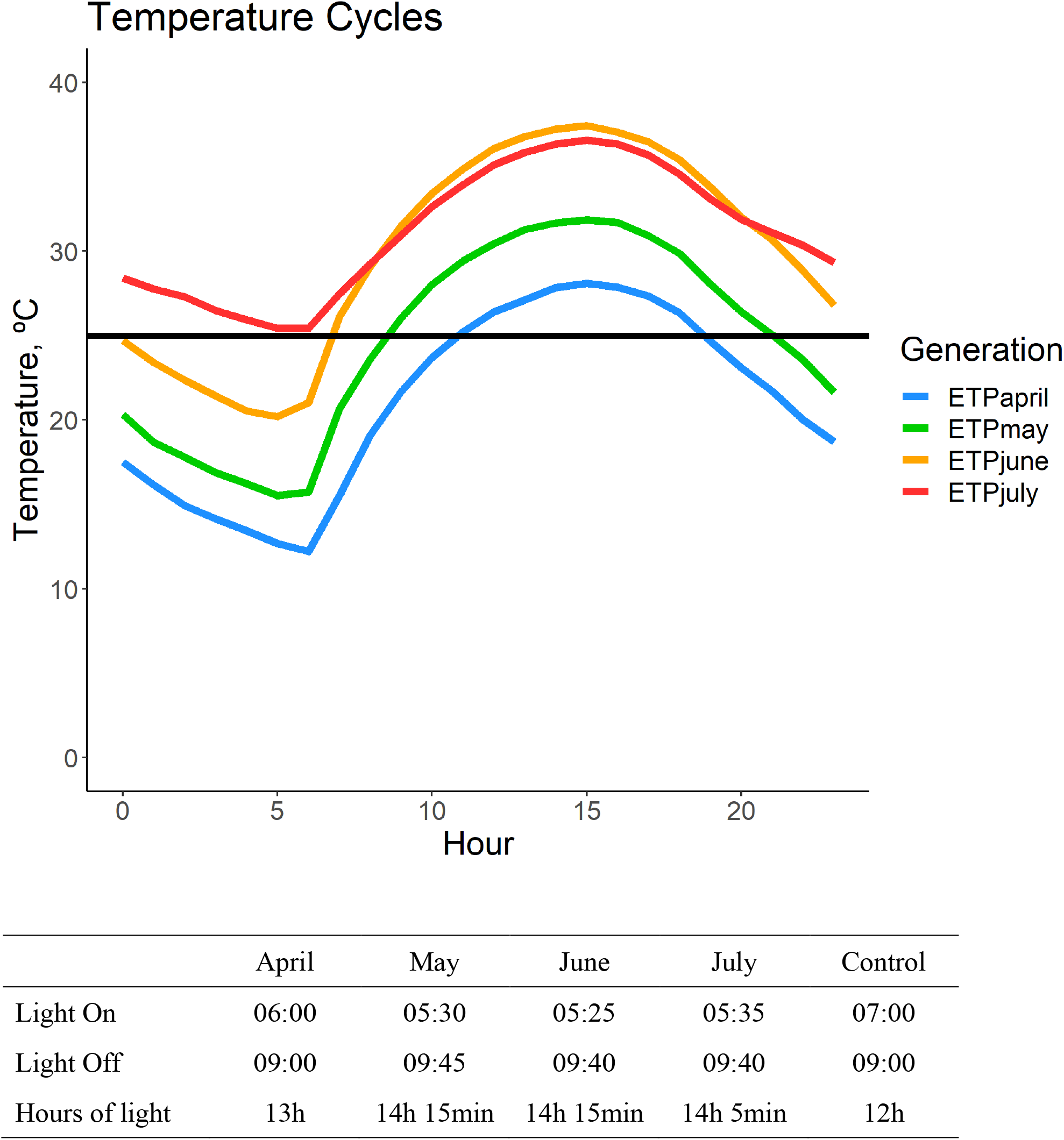
Experimental temperature cycles shown for each of the temperature treatments used. Solid black line denotes the constant 25°C temperature cycle used for the CON population. Hour 0 = 12:00 AM. Box below indicates the light regime for each cycle.

**Fig 2.**
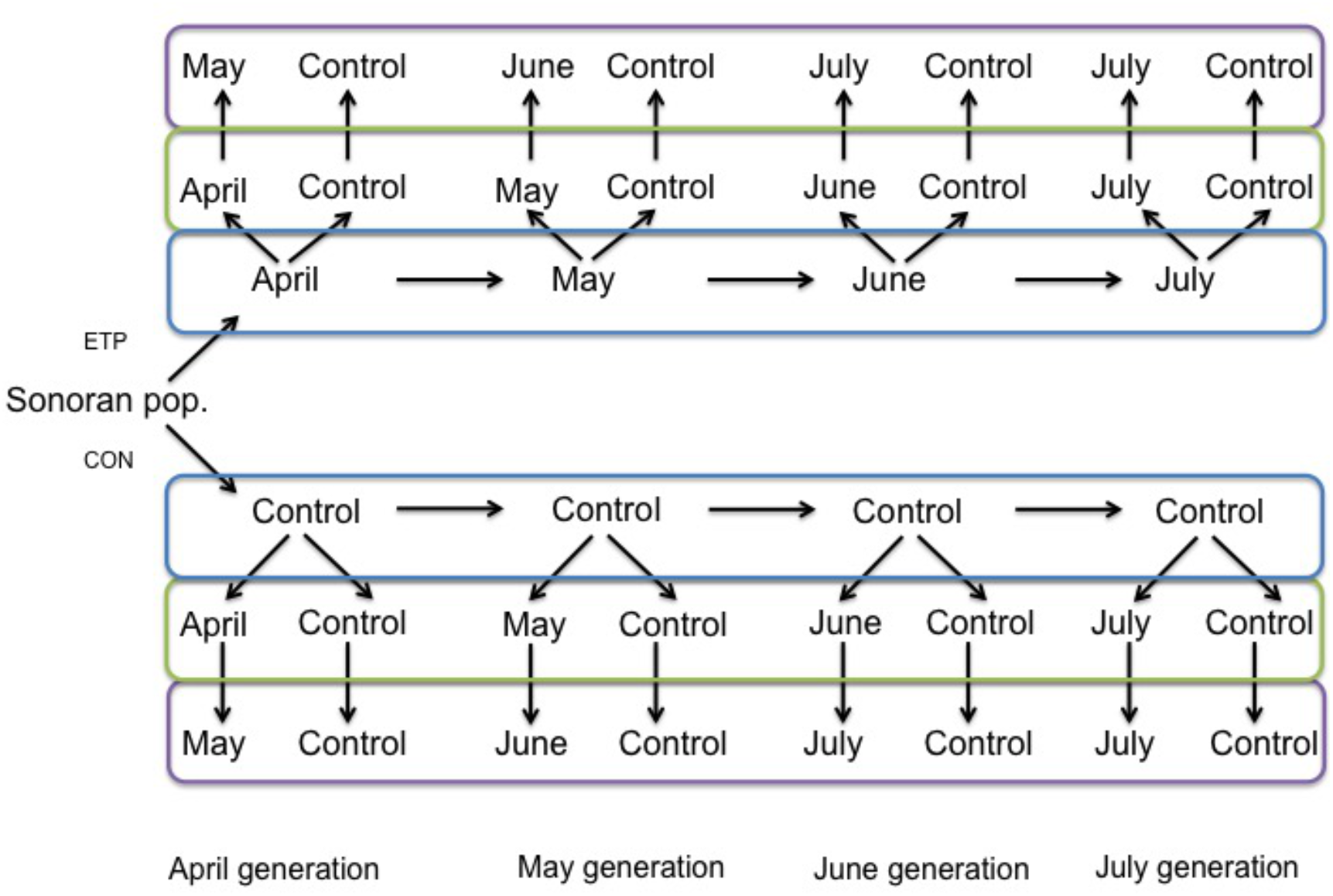
Experimental setup used in this study. Sonoran population was divided into the experimental temperature population (ETP) and the constant 25°C population (CON). Horizontal arrows represent offspring movement, while other arrows represent adult movement. Generations shown in the blue box represent the *Developmental* temperature used, while the green and purple boxes represent the *Test* temperatures used. Phenotypes adult heat stress, larval heat stress, and metabolic pools were carried out at the blue box level and have no *Test* factor as all experimenting was carried out directly from the *Developmental* temperature. Adult activity, fecundity, developmental time, viability, desiccation, and nutritional stress resistance were carried out at the level of the green box. Longevity assay was carried out at the level of the purple box, with adults moving through the green level for testing fecundity first.

### 2.4 Adult activity level assays

Each generation, flies were collected from the pooled cages on the 11^th^ day (11-16 days posteclosion adults). After light CO_2_ anesthesia, individual flies were placed in individual small glass vials (50 mm long, 6 mm outside diameter, 5 mm inside diameter) with a wet cotton ball. A total of 20 females and 20 males from each population (CON and ETP) were collected for observation in both a 25°C constant chamber (14:10 light:dark) and the given temperature cycle being tested at that generation. Flies were tested by population, with vials laying horizontally in two columns of females flies next to two columns of male flies. Flies were recorded for a 24-hour period and filmed using a Raspberry pi camera. Ethoscope was used to identify the position of the fly per frame (Geissmann et al., 2017) at a rate of one frame per second. Movement was recorded as the distance between each position per frame and was binned into hour long segments.

### 2.5 Egg-to-adult viability and adult life history assays

As described above, each generation, flies were collected from the pooled cages on the 11^th^ day. Females were put into banana-molasses food 30 ml vials sprinkled with yeast in groups of five. Five replicate groups were tested in both temperature cycles (e.g. 25°C and May) from both the CON and ETP. Each day for 10 days the flies were transferred into a new banana-molasses food vial and eggs oviposited in the food media were counted visually once the flies were removed. These eggs were maintained in the temperature cycle the adults were tested in until eclosion, the date of which was recorded to measure both developmental time and egg-to-adult viability.

Following the 10 days of the fecundity assay, the females whose eggs were counted were pooled together in a banana-molasses food vial and put into the next temperature cycle-e.g.-flies whose fecundity assay was done in April went into the May temperature cycles. This change was done to maintain consistency with temperatures flies would experience in the wild. Flies tested in the 25°C temperature chamber remained at 25°C. The flies were observed daily until their death. Each female group had a matched male group.

### 2.6 Adult and larval heat stress tolerance

For adult heat stress, each generation, 24 virgin females and males were sampled per population for the heat stress assay. Virgin flies for the heat assay were collected from the eclosion vials after the first five days of collection for the cages (see above). Collected virgins were used at 3 days of age. Heat stress began around 8 AM when the cycling temperatures are relatively low (see Fig. 1). Heat stress was carried out in a circulating water bath with the temperature controlled by a Thermo Scientific Circulator (AC 200). Heat stress temperatures began at 30°C and increased at a constant rate for the first hour and a half to 40°C. Flies were then held in the 40°C circulating water bath until they no longer respond to light. Single flies were held in completely empty 1-dram glass vials arranged randomly in an acrylic frame and checked continually; time was recorded when the fly was determined dead by: no movement of any limb/wing when stimulated by a bright LED white light and after tapping of the container. Flies were checked every two minutes after finishing the previous check.

Larval heat tolerance was also measured each generation. Following the monthly egg collection, a new egg collecting plate was placed in each population’s cage. The egg plate was removed from the cage after ∼20 hours. The eggs were then left in their respective temperature cycles to develop for about 24 hours. At that time, first instar larvae were picked from the plates and put into banana-molasses food vials in groups of 30. The vials were then placed into the circulating water bath, as described above. Temperatures began at 30°C and increased at a constant rate for the first hour and a half to 40°C. For the following two hours the larvae were held at 40°C. Vials with larvae were then placed back into their respective temperature cycle chambers, monitored and the daily number of eclosed adults per vial recorded.

### 2.7 Desiccation and nutritional stress tolerance

Following both the June and July generations, a desiccation assay was performed on the flies. In the June generation, the desiccation assay was carried out with both populations being assayed at 25°C. In the July generation, both populations were assayed both at 25°C, as well as in the July temperature cycle. Flies were collected about 15 days after being put into the pooled cages, making them 15-20 days old. Flies were lightly anesthetized using CO_2_, then placed individually into the bottom of clean 30 ml glass vials. Next, half of a cotton ball was placed about 2 cm above the bottom of the vial. Approximately 5 ml of indicating Drierite desiccant (W.A. Hammond) was put on top of the cotton ball, then the vial was sealed using parafilm (Bemis). A total of 20 males and 20 females per population per temperature cycle were used. The flies were checked every 2 hours for movement. Flies were recorded as dead once they underwent two subsequent checks without any movement.

Following the July generation, a low nutrition assay was performed for both populations at both 25°C and the July temperature cycle. Flies were collected about 15 days after being put into the pooled cages (15-20 days old). Flies were lightly anesthetized using CO_2_, then placed into vials with low nutrient media. The media was 1% agar (14.8 mg agar in 1 L of DI water) with 2.22 g methylparaben dissolved in 22 mL of 95% ethanol, providing them with a small source of nutrition. Per population and temperature cycle, 20 males and 20 females were used. The vials were closed using a condensed cotton plug (Flugs, Genesee Scientific) and placed in the chambers and their respective cycle temperatures. The flies were checked every 6 hours for movement. Flies were recorded as dead once they underwent two subsequent checks without any movement.

### 2.8 Metabolic pools content

Each generation, five replicates of five flies each per sex were sampled from the CON and from the ETP. These adult flies (10-15 days old) were frozen at -20°C for approximately one hour, then weighed on a Mettler Toledo XS3DU microbalance. The samples were placed in an incubator for 24 hours at 50°C and then reweighed. Dried flies were homogenized using a powered hand pestle in 300 µl of phosphate buffer (25 mmol/L KHPO4, pH 7.4). Following initial homogenization, 700 µl of buffer was added, mixed by vortexing and 950 µl of supernatant was saved for the metabolic pool assays. Glycogen assays (Pointe Scientific G7521-500, Roche 10102857001, Thermo Sci J16445-06), triglyceride assays (Pointe Scientific T7532-120, T7531-STD) and soluble protein assays (Sigma B9643-1L) were then carried out in triplicate according to the kit instructions. The means were normalized using the respective sample dry weights.

### 2.9 Data analysis

All data analysis was carried out in R 3.3.3 (R Core Team, 2013). The complete set of output tables of all analyses, including pairwise tests, are publicly available at OSF (https://osf.io/eqbgk).

#### 2.9.1 Horizontal test

In order to address generational differences within the ETP or CON populations (ie. across the generations, see Fig. 2 - comparisons within a blue box), the following general test was used for all phenotypes:

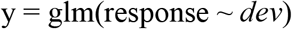

Where *dev* is the *developmental* temperature cycle in which the group was raised (April, May, June or July). This test allows me to address how the phenotypes have changed as we move from the early generations to the late generations. At the same time, we could compare the CON developing in constant 25°C, allowing us to identify changes that originate from sources other than temperature/photoperiod changes throughout this experiment. In doing so, we could identify if there were changes with the ramp up of the summer conditions.

#### 2.9.2 Vertical test

In order to address differences between the populations, but within a single generation (see Fig. 2, comparisons in a column), the following general test was used for all phenotypes:

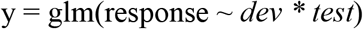

Where *dev* is the *developmental* temperature cycle in which the group was raised, and *test* is the temperature cycle in which the phenotype was measured. The use of these tests allowed us to uncover any phenotypic advantage when tested in the same temperature cycle in which the population developed.

#### 2.9.3 Analysis of individual phenotypes

Activity data was analyzed with all individuals per treatment pooled together and the data split into four time blocks. Each block is composed of either the first or second half of the light and dark cycles. Average activity was estimated for each block for each treatment. Data was log transformed for normality and both the *horizontal* and *vertical* tests were performed. *Sex* was included as a fixed variable, and the hour of the day (*hour*) and the individual being assayed (*ID*) were included as random factors in the lmer() call (Bates et al., 2015).

Fecundity data was analyzed with all individuals per replicate pooled together. The 10 days over which egg counting was done was blocked into an early (days 1-5) and late (days 6-10) block. Both the *horizontal* and *vertical* tests were performed with glmer() and a poisson family link function. The early/late *blocks* were included as a fixed variable and each replicate was included as a random factor. Additionally, fecundity was analyzed with the early/late blocks pooled together. For this analysis, *horizontal* and *vertical* tests were performed with glmer() and a poisson family link function, with replicate included as a random factor.

Developmental time was recorded as the time, in days, from oviposition to eclosion. A glmer() with a poisson family link function was used, and replicate was included as a random factor. Both the *horizontal* and *vertical* tests were performed. For this phenotype, in the *vertical* test, *dev* represents the developmental temperature cycle of the parental generation of the generation being tested, and *test* represents the temperature cycle of the individuals being reported.

Viability was calculated from the number of eggs that developed to adulthood, and the number of eggs that did not. Both the *horizontal* and *vertical* tests were performed. In this phenotype, in the vertical test, *dev* represents the developmental temperature cycle of the parental generation of the one being tested, and *test* represents the temperature cycle of the individuals being reported. A glmer() with a binomial family link function was used, and replicate, number of eggs and the day of laying were included as random factors.

Longevity was analyzed using a coxme() model using the number of days that individual flies lived for (Ripatti and Palmgren, 2002). *Sex* was included as a fixed effect, and the date that the experiment began was included as a random factor. Both *horizontal* and *vertical* tests were performed.

Adult heat stressed was based on the minutes that individuals survived during the heat stress experiment. A coxph() model was used with *sex* included as a fixed effect. Both horizontal and vertical tests were performed. Larval heat stress was standardized based on a control group that did not undergo heat stress and a Gaussian family, identity link was used with a glm() model for its analysis. Both *vertical* and *horizontal* tests were performed.

Both phenotypes were analyzed using the number of hours it took the flies to die under the respective stress. The tests were done with using the coxph() model, and included *sex* as a fixed effect. Only *vertical* tests were performed on these phenotypes, as not all generations were tested for this phenotype.

Glycogen, triglyceride, and protein contents were collected as a ratio of the total mass. Thus for analysis, the arcsine square root of the content in nanograms was used with an lm() which included *sex* as a fixed effect. Both the *horizontal* and *vertical* tests were performed. For dry mass, an lm() model with *sex* as a fixed effect was used in both the *horizontal* and *vertical* tests.

## 3. Results

### 3.1 Field temperatures

As shown in Fig. 3 temperatures inside necrotic cactus are lower than those on the exterior of the cactus (on average, maximum temperatures in the cactus necroses are 6% lower). Across the period recorded (May to October) the minimum and maximum temperatures recorded for the external and the internal probes were, 8.2°C /54.1°C and 9.0°C /50.7°C, respectively.

**Fig 3.**
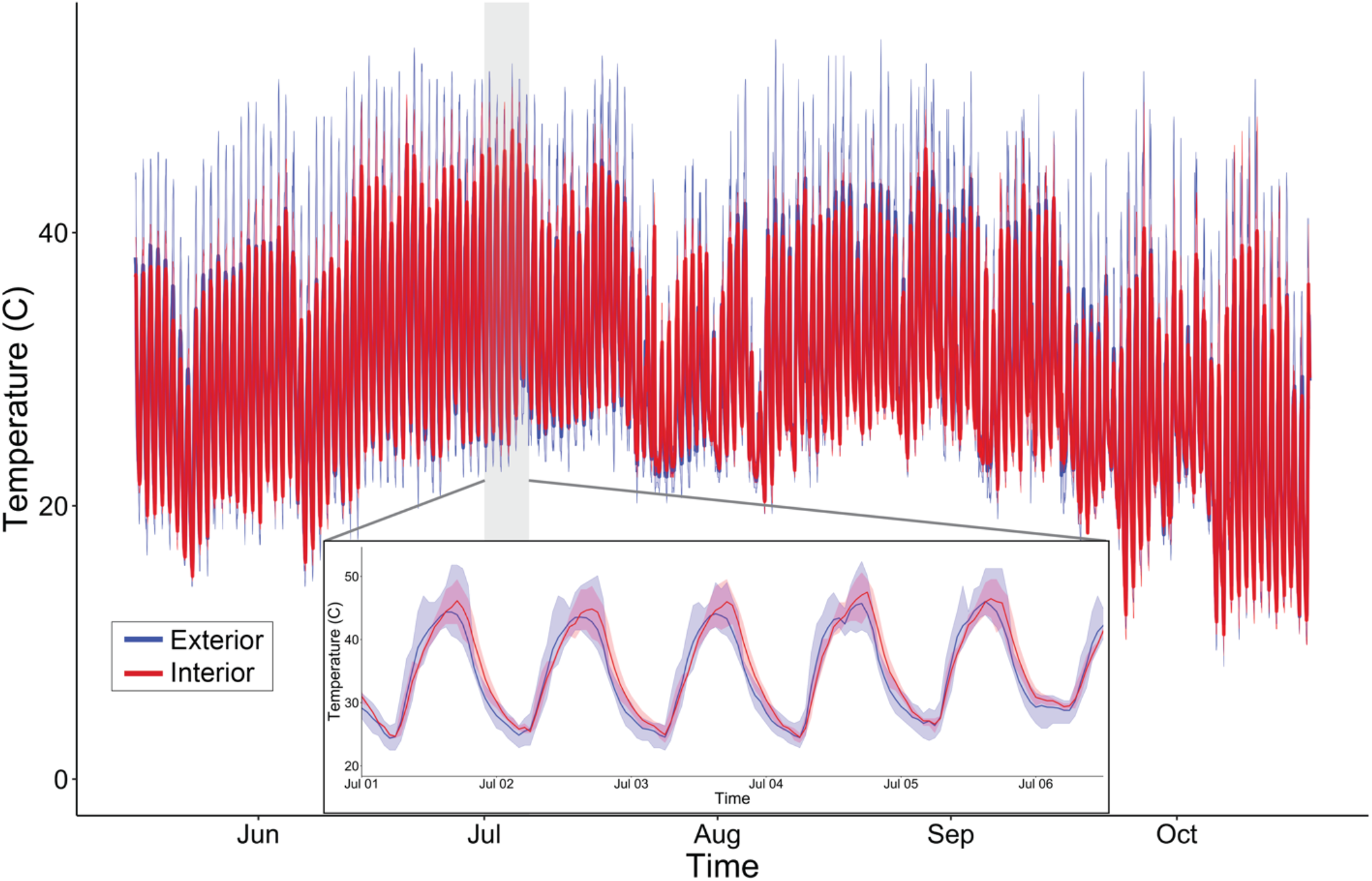
External (blue) and internal (red) cactus probe temperatures in Organ Pipe National Monument, Arizona (May - October 2007). Line indicates mean temperature and shaded area reflects the maximum and minimum temperatures recorded. Temperature profile of gray area (shown in insert) illustrates measurements for 5 days in July.

### 3.2 Adult activity

Adult activity patterns for populations and treatments are shown in Fig. 4. Analysis performed on activity data of flies from the ETP, with the *horizontal* test showed a significant *dev* effect (*χ*^*2*^ = 75.2, P < 0.001). Tukey’s *post hoc* testing showed significant differences among all pairwise comparisons (P < 0.001) of the developmental temperatures, except for between May and July (P = 0.08). In the *horizontal* test of the CON, Adult activity data showed no significant *dev* effect, however, it did show a significant *dev* by *sex* interaction (*χ*^*2*^ = 13.1, P = 0.004). However, Tukey’s *post hoc* test showed no significant differences between any of the pairwise comparisons, suggesting that the phenotype was very similar across the generations of the CON.

**Fig 4.**
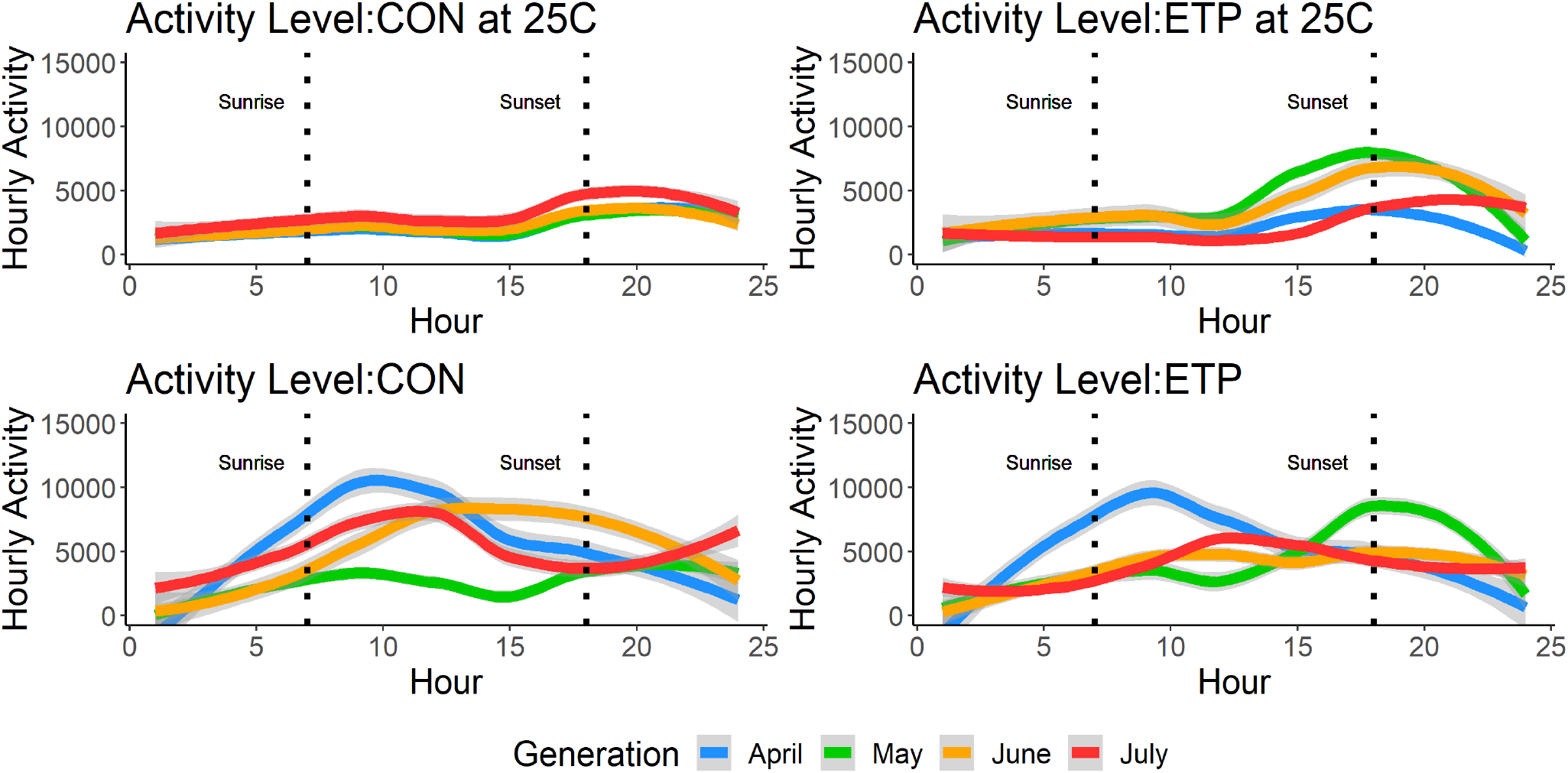
Mean adult activity across a 24-hour cycle in all populations and treatment. Hour zero is midnight. CON in 25 is CON tested at constant 25°C. CON is CON tested in the experimental temperature cycle that corresponds to the given generation, the same notation is used for the ETP. The dotted lines denote the average times of the sunrise (left) and sunset (right) across all temperature cycles. The gray lines indicate 95% confidence interval.

For the *vertical* tests, comparisons between ETP_april_ to CON_april_, and ETP_may_ to CON_may_ both showed a significant effect based on the *test* (*χ*^*2*^= 9.59, P = 0.001 and *χ* ^*2*^ = 38.4, P < 0.001, respectively), although neither showed a significant *dev* effect. The same trend was seen in the July generation (*χ*^*2*^= 14.6, P = 0.001). For the June generation however, both the *dev* (*χ*^*2*^= 7.91, P = 0.005) and the *test* (*χ*^*2*^= 22.6, P < 0.001) effects were significant. The recovery of the phenotype seen in the constant 25°C by the ETP shows the presence of phenotypic plasticity for this trait.

### 3.3 Life history assays

#### 3.3.1 Fecundity

Fecundity, measured as the number of eggs laid per day per five females, was analyzed as two blocks - the first five days of laying, termed the “early” block, and the last five days of laying, termed the “late” block (Fig. S1). When looking at the *horizontal* tests of the ETP, there is a significant *dev* by *block* interaction (*χ*^*2*^ = 513.9, P < 0.001). The Tukey’s *post hoc* test shows that all comparisons were significant (P < 0.05), except May_early_:July_early_, May_late_:June_late_, and June_late_:July_late_. highlighting the differences between ETP_april_ and the other generations. Likewise, in the *horizontal* test for the CON, the fecundity data also showed a significant *dev* by *block* interaction (*χ*^*2*^ = 1225.38, P < 0.001). The ETP the Tukey’s *post hoc* tests for the CON revealed non-significant pairwise tests between April_early_:June_late_ and April_early_:July_late_, the rest of the comparisons were all significant (P < 0.05). The high level of variability across the CON in constant 25°C highlights generational (i.e. environmental) differences that exist regardless of temperature for this experiment. Likewise, the comparisons across populations, between generations, display high levels of variation. On average, the ETP produced the greatest number of eggs per day in the June generation, regardless of *test* temperature cycle (Fig. 5).

**Fig 5.**
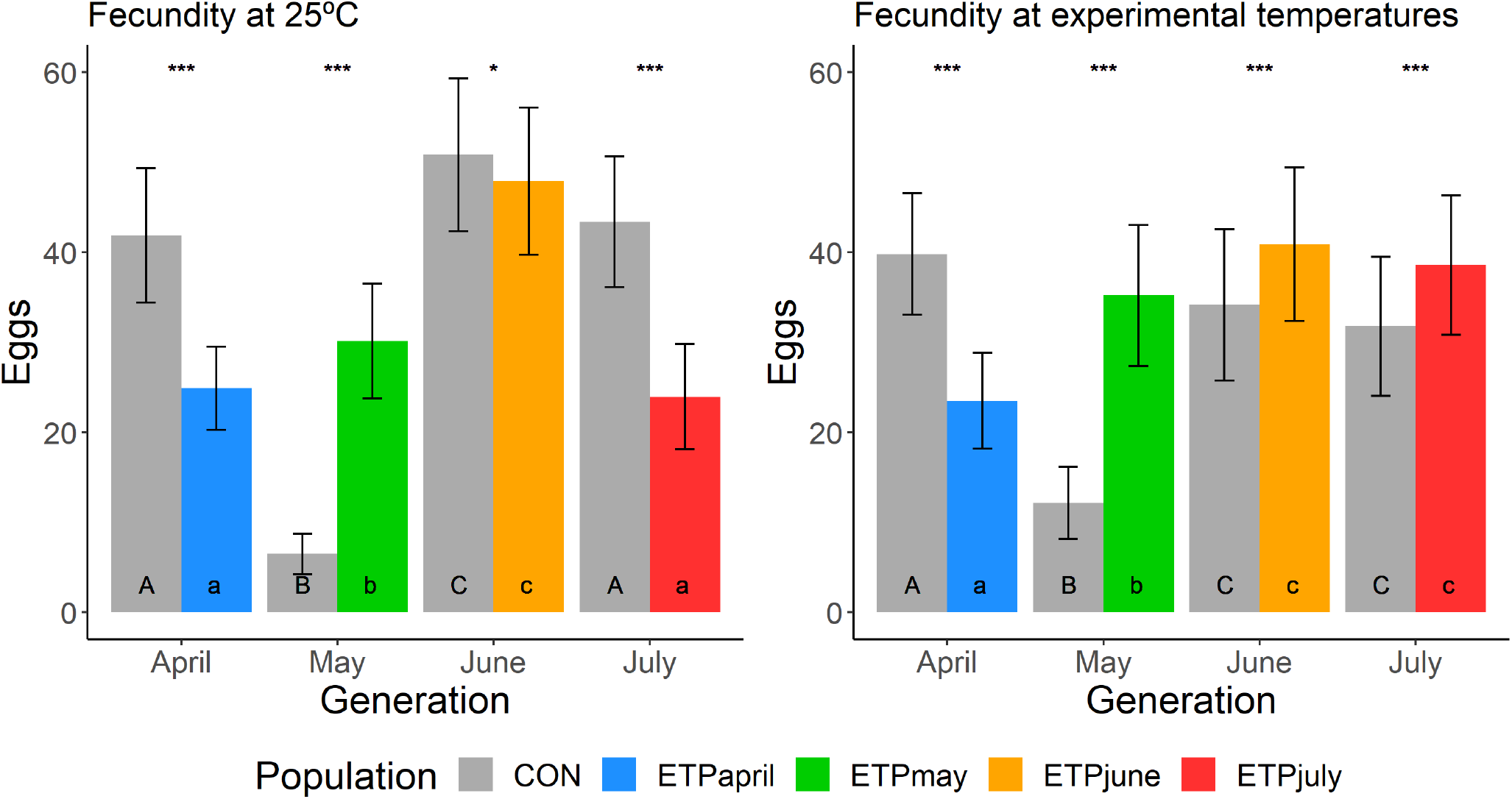
Mean (± std. error) of egg production (number of eggs laid per day per five females) across treatments. Asterisks indicates significance (*p < = 0.05, **p < = 0.01, ***p < = 0.001) between CON and ETP populations of the same generation. Uppercase letters indicate significant differences across generations of the CON population, while lowercase letters indicate significant differences across generations of the ETP.

#### 3.3.2 Developmental time

Developmental time was measured for the eggs laid during the fecundity assay (Fig. S2). For the *horizontal* test of the ETP, there was a significant effect of *dev* (*χ*^*2*^ = 391.8, P < 0.001). All pairwise tests between generations were significant (Tukey’s *post hoc*, P < 0.001). The average development time decreased from 20 days in the ETP_april_ to 13 days in ETP_july_. In the *horizontal* test of the CON, *dev* was also significant (*χ*^*2*^ = 104.0, P < 0.001). *Post hoc* analysis showed that the CON_june_ was different than all other generations (P < 0.001), displaying a slightly longer development time than the others.

For the *vertical* tests, in the April generation, there is no significant difference between the two populations (ETP_april_ and CON_april_), although when they are both tested in the constant 25°C, the two populations show slightly different phenotypes (*χ*^*2*^ =5.59, P = 0.018). Both ETP_may_ and CON_may_ performed the same when tested in their respective temperature cycle, so only *test* was significant (*χ*^*2*^ = 4.28, P = 0.038). In the June generation, the ETP_june_ and CON_june_ performed differently from each other when tested in both the June temperature cycle (*χ*^*2*^ = 15.18, P < .001) as well as in the constant 25°C (*χ*^*2*^ = 197.8, P < .001), with the CON displaying the longer average development time in both temperature regimes. Lastly, ETP_july_ and CON_july_ performed differently from each other regardless of which temperature cycle they were tested in (*χ*^*2*^ = 103.2, P < 0.001), although ETP_july_ when tested in the July temperature and the constant 25°C, were not significantly different.

#### 3.3.3 Egg-to-adult viability

Viability was measured by comparing the number of eggs laid for the fecundity phenotype to the number of adults that emerged from those eggs (Fig. S3). In the horizontal test of the ETP, *dev* (*χ*^*2*^ = 222.6, P < 0.001) was significant. Viability increased from 30% in ETP_april_ to 50% in ETP_may_, but then dropped down to 15% in ETP_june_ and even lower to 7% in ETP_july_. Tukey’s *post hoc* tests showed all comparisons to be significant (P < 0.001). In the *horizontal* test of the CON, *dev* (*χ*^*2*^ = 101.8, P < 0.001) was again significant, but the trend over the generations was very different from that of the ETP. The CON generations displayed a steady increase in viability each generation, from 24% in CON_april_ to 45% in CON_july_. CON_april_ and CON_may_ were not significantly different, nor were CON_june_ and CON_july_.

In the *vertical* tests, the April generation showed a significant *dev* by *test* (*χ*^*2*^ = 6.25, P = 0.012) effect. CON_april_ performed the same in both temperature cycles, but ETP_april_ performed better in the experimental temperature cycle. When performance was compared in the same temperature cycles, the two populations did not differ in performance. In the May generation, only the *dev* effect (*χ*^*2*^ = 128.2, P < 0.001) was significant. Viability was higher in ETP_may_ than in CON_may_, regardless of the temperature cycle they were tested in. In the June generation, there was a significant *dev* by *test* effect (*χ*^*2*^ = 11.4, P < 0.001). Both ETP_june_ and CON_june_ had lower viability in the experimental temperature cycle than in the constant 25°C, but ETP_june_ had higher viability than CON_june_ in the experimental temperature cycle, while CONj_une_ had higher viability than ETP_june_ in the constant 25°C. In the July generation, the same general trend as in the June generation was observed.

#### 3.3.4 Longevity

Longevity was measured on the exact same flies used for the fecundity assay; thus, flies were already aged 20-25 days when the longevity testing begun. ANOVA tests on the longevity data in the horizontal tests on ETP showed significant *dev* by *sex* interaction (*χ*^*2*^ = 17.8, P < 0.001). Pairwise comparisons of flies across generations showed that ETP_may_, ETP_june_ and ETP_july_ males had a similar longevity, but only ETP_may_ and ETP_july_ females were similar. Overall, averages across generations ranged from the longest in the April generation at 26 days, to the shortest in the May generation at 3 days (Fig. 6).

**Fig 6.**
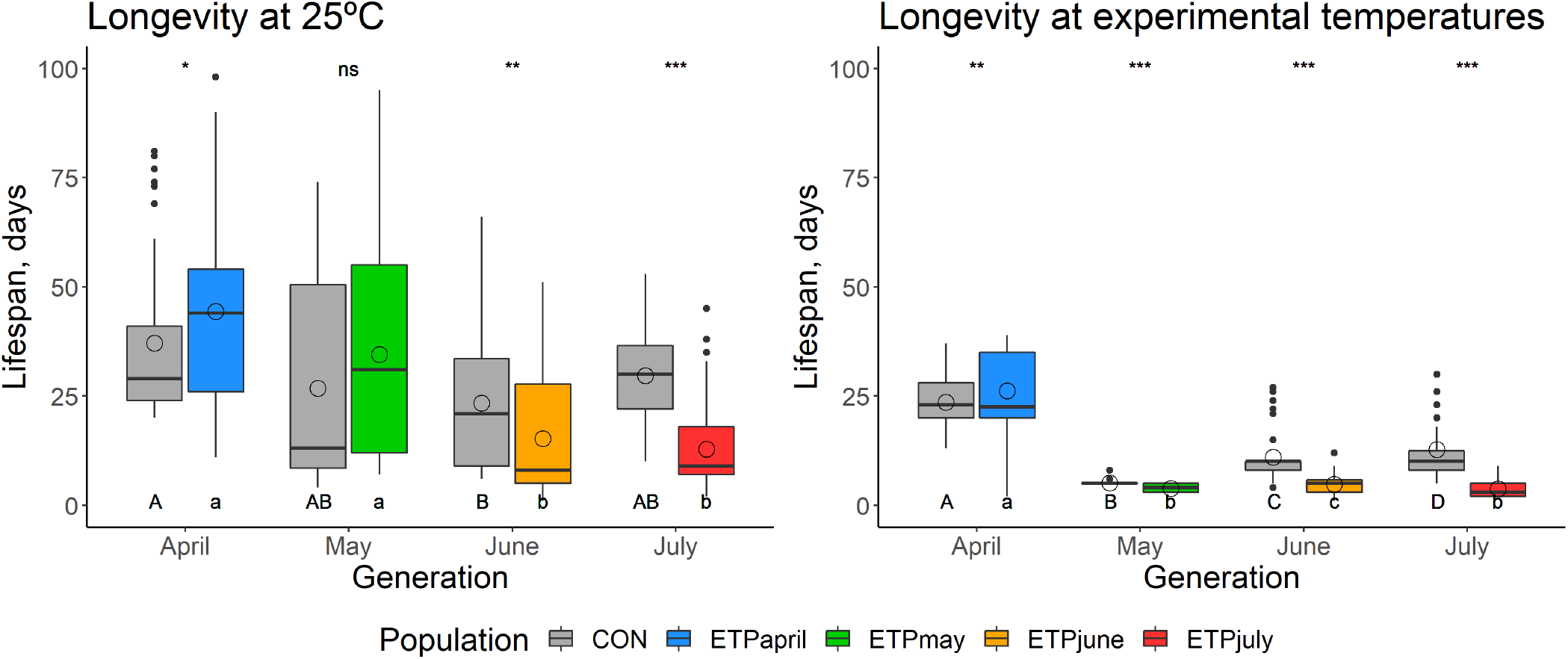
Box plot of longevity, in days, of individuals surviving after completing the fecundity assay (see text). Box plot horizontal lines indicate the first, second, and third quartiles and whiskers show the extreme upper and lower values within 1.5 times the interquartile range (IQR). Solid circles are values beyond 1.5× IQR. Open circles denote the average value for the group. Asterisks indicates significance (*p < = 0.05, **p < = 0.01, ***p < = 0.001) between CON and ETP populations of the same generation. Uppercase letters indicate significant differences across generations of the CON population, while lowercase letters indicate significant differences across generations of the ETP.

For the *horizontal* test of the CON, a significant *dev* by *sex* interaction was observed (*χ*^*2*^= 9.0, P = 0.029), however, the Tukey’s *post hoc* test showed that the differences came mainly from differences in females, as there were no significant differences between males of different generations. Significant differences did appear between females of April and June, April and May and June and July (Tukey’s *post hoc*, P < 0.001, P = 0.001, P = 0.019, respectively). Averages for the CON ranged from the longest in the April generation at 37 days and the shortest in the June generation at 23 days - a much smaller range than in the ETP.

For the *vertical* tests, in the April generation, we see differences based on the testing temperatures (*χ*^*2*^ = 9.926, P =0.001), as well as an effect of *dev* by *sex* interaction (*χ*^*2*^ = 12.548, P < 0.001). In the May generation, there is a significant *dev* by *test* by *sex* interaction (*χ*^*2*^ = 9.183, P = 0.002). The June generation showed differences in both the ETP males and females (Tukey’s *post hoc*, P < 0.001), as well as in the CON males and females (Tukey’s *post hoc*, P = 0.016). The July generation instead showed that the males and females within a population and a testing temperature cycle were the same (Tukey’s *post hoc*, P = 0.81 in ETP, P = 0.87 in CON). Overall, the ETP and the CON tested in the same temperature cycle showed similar trends and ranges of average longevity.

### 3.4 Adult and larval heat stress

For *vertical* tests of the adult heat stress, both populations had a significant *dev* by *sex* effect (*χ*^*2*^ = 8.55, P = 0.035 and *χ*^*2*^ = 11.7, P = 0.008, for ETP and CON respectively). In both populations, the average female survival time was about 26 minutes longer than the average male survival time, except for the case of the CON_july_, where the male survival time was 26 minutes higher than the female survival time (Fig. 7). Across the ETP, all Tukey’s *post hoc* comparisons were significantly different (P < 0.05), excluding those involving June and July. Each generation had a subsequently equal or longer survival time than the generation before it. In the CON, on the other hand, CON_may_ was significantly different from all other generations (the average being about 25 minutes less than the other generations), but the rest of the generations were equivalent. When comparing the two populations in the vertical tests, we see a significant *dev* effect in all comparisons. Only did the CON_april_ survive longer than the ETP_april_; in all other generations, the ETP survived longer.

**Fig 7.**
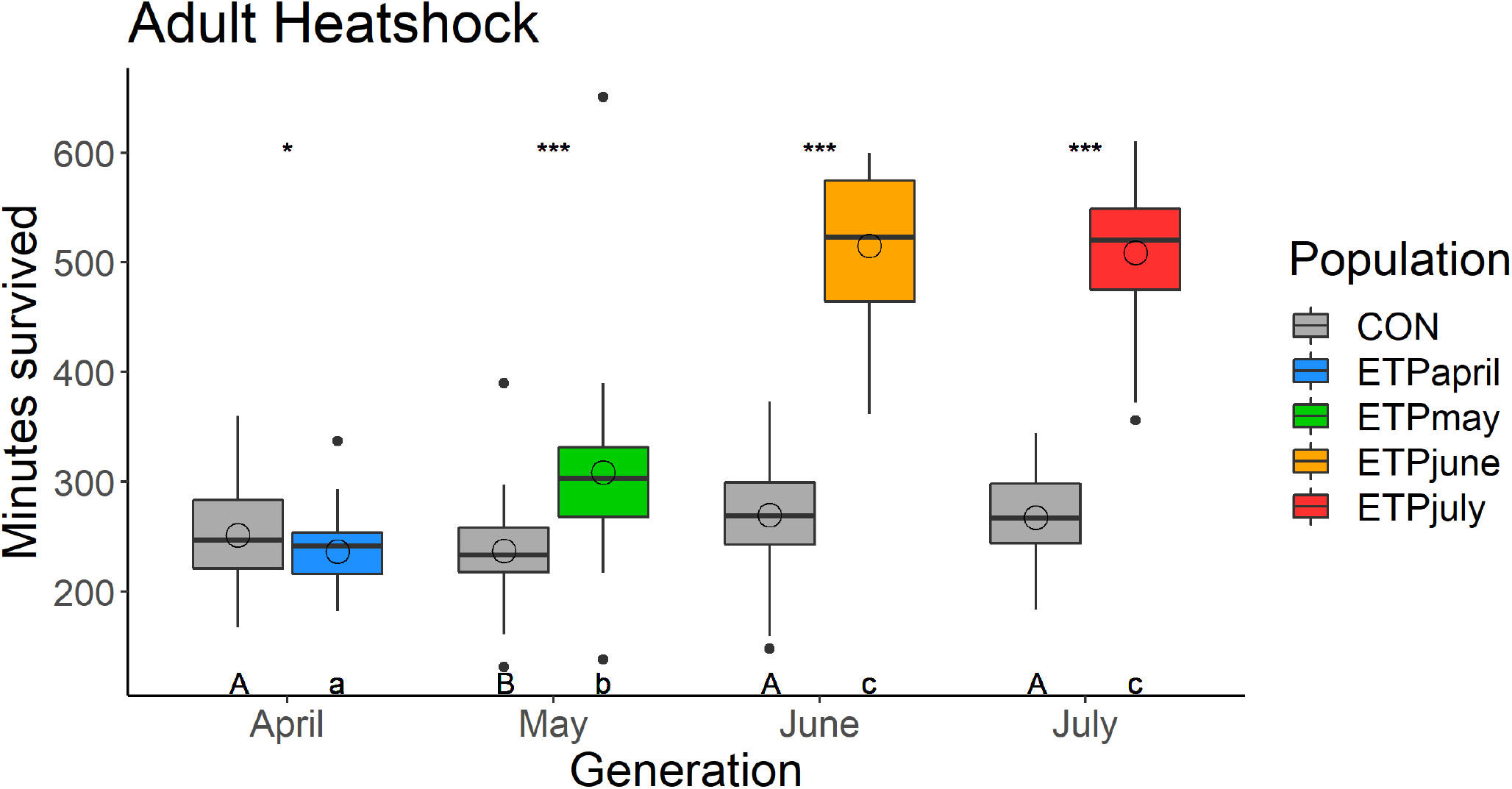
Adult heat stress measured as the time (minutes) to non-responsiveness in the circulating water bath. Box plot horizontal lines indicate the first, second, and third quartiles and whiskers show the extreme upper and lower values within 1.5 times the interquartile range (IQR). Solid circles are values beyond 1.5× IQR. The open circle denotes the average value for the group. Asterisks indicates significance (*p < = 0.05, **p < = 0.01, ***p < = 0.001) between CON and ETP populations of the same generation. Uppercase letters indicate significant differences across generations of the CON population, while lowercase letters indicate significant differences across generations of the ETP.

For the horizontal test of the ETP, larval heat stress had a significant *dev* effect (*χ*^*2*^ = 96.8, P < 0.001). ETP_april_ was significantly different from all other pairwise comparisons. The survival of offspring increases from a standardized 0.2% in the April generation, to as high as 86% and 68% in June and July, respectively. For the *horizontal* test within the CON, there was also a significant *dev* effect (*χ*^*2*^ = 15.1, P = 0.002). The average standardized survival of the 25°C was 15% in all generations except for the June generation where it dropped to 5%. In the *vertical* tests, there was a significant *dev* effect in all generations (April, *χ*^*2*^ = 29.1, P < 0.001; May, *χ*^*2*^ = 77.5, P < 0.001; June, *χ*^*2*^ = 49.705, P < 0.001; July, *χ*^*2*^ = 103.9, P < 0.001). The ETP outperformed the CON in every generation, aside from CON_april,_ which had a higher survival than ETP_april_ (Fig. 8).

**Fig 8.**
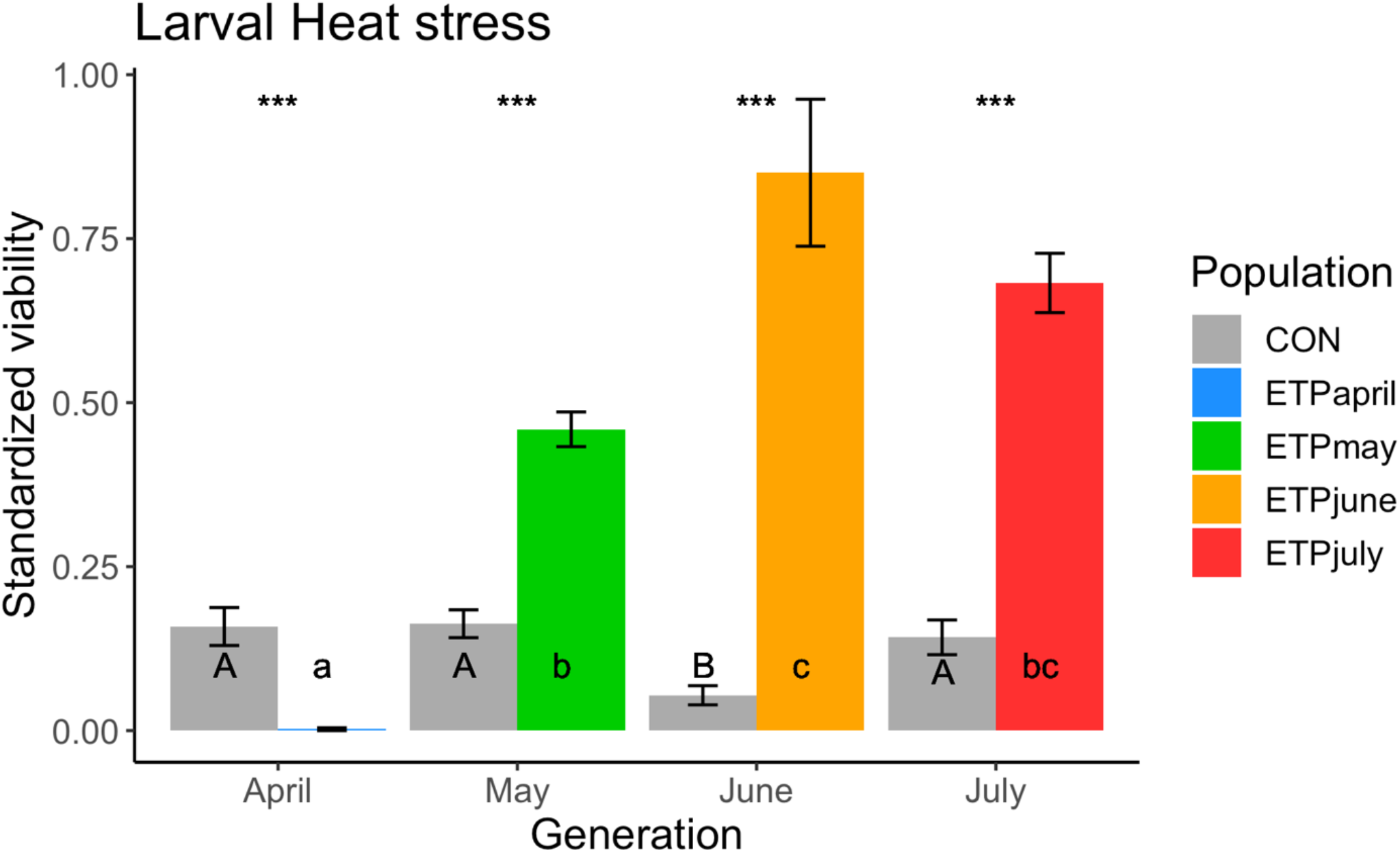
Larval viability (± std. error) following heat stress standardized by the survival of the non-heat stressed group. Asterisks indicates significance (*p < 0.05, **p < = 0.01, ***p < = 0.001) between CON and ETP populations of the same generation. Uppercase letters indicate significant differences across generations of the CON population, while lowercase letters indicate significant differences across generations of the ETP.

### 3.5 Desiccation and nutritional stress

Desiccation resistance was measured for both populations only in June and July generations. For the June generation, populations were only tested in the constant 25°C, while for the July generation, the populations were tested in both the constant 25°C and the July temperature cycle (Fig. S4). For the June generation, there was a significant *dev* by *sex* interaction (*χ*^*2*^ = 4.23, P = 0.039). *Post hoc* tests revealed that males and females from the same population were not different, but all other pairwise comparisons were significantly different (Tukey’s *post hoc*, P < 0.001). In the July generation, both the *test* (*χ*^*2*^ = 82.5, P < 0.001) and the *dev* by *sex* interaction (*χ*^*2*^ = 15.4, P < 0.001) were significant. In all three testing temperature cycles used, the ETP lived longer than the CON.

The nutritional stress assay was only measured in the July generation, but in both temperature cycles (Fig. S5). The *test* effect (*χ*^*2*^ = 107.2, P < 0.001) and the *dev* by *sex* interaction (*χ*^*2*^ = 10.2, P < 0.001) were both significant. Both populations lived longer when tested in the constant 25°C than when they were tested in the July temperature cycle, and the CON lived longer than the ETP, regardless of which temperature cycle they were tested in.

### 3.6 Metabolic pools content

Dry fly mass was measured to standardize metabolic pool content across samples. In both populations and all generations, the average female weight was larger than the average male weight (Fig. S6). Weights varied significantly across the generations for both populations (*F* = 89.3, P < 0.001 and *F* = 41.7, P < 0.001, for ETP and CON respectively). The ETPs showed a trend of decreasing across all generations from 2.7 mg in ETP_april_ to 1.6 mg in ETP_july_. The CON showed decreasing trends from 2.7 mg CON_april_ to 2.1 mg in CON_june_, but CON_july_ increased slightly to 2.3 mg, in stark contrast to ETP_july_. When doing the vertical tests, there is no effect of *dev* on the weight, except for the July generation (*F* = 66.1, P < 0.001). In ETP_july_, the weight drops abruptly from the averages of the other population and other generations.

Glycogen content was measured as the proportion of the mass of which is made up of glycogen. For the horizontal tests, within the ETP, glycogen had a significant *dev* by *sex* interaction (*F* = 3.17, P = 0.037). In the ETP, glycogen content was higher in the males than in the females, except for in ETP_july_, where females had higher glycogen content (Fig. S6). The ETP_april_, ETP_may_, and ETP_june_ were not significantly different from one another; however, the ETP_july_ had a lower glycogen content than the rest of the generations (Tukey’s *post hoc*, P < 0.01 for all comparisons). In the CON, the females had lower average glycogen content than the males in all generations. The glycogen content in the CON varied across the generations, but in all cases except for the July generation, the CON had a lower glycogen content than the ETP. This difference was driven by the ETP_July_ males who have lower glycogen content than both the ETP_july_ females (though not significantly), and the CON_july_ males (Tukey’s *post hoc*, P = 0.005).

Triglyceride content was measured as the proportion of the mass that consists of triglyceride. Triglyceride content in both populations was higher in all generations in females than in males. In the ETP, there was a significant *dev* effect (*F* = 4.68, P = 0.008) (Fig. S6). ETP_july_ was different from both ETP_may_ and ETP_june_, with a lower average triglyceride content in ETP_july_ than any other generation. The CON likewise had a significant *dev* effect (*F* = 6.08, P = 0.002). The CON_may_ had lower triglyceride content than either the CON_june_ or CON_July_ (P < 0.001 and P = 0.004, respectively). In the vertical tests, we see no *dev* effect in the May generation, but all other generations have a *dev* effect (Table S18). July is the only generation with a significant *dev* by *sex* effect (*F* = 4.86, P = 0.04), and the Tukey’s *post hoc* analysis show that this is driven again by low triglyceride content in the ETP_july_ males (P < 0.05).

Protein content was measured as the proportion of the mass that is made up of protein. In the ETP, males had higher protein content than females in all generations except for the ETP_may_, where females’ protein content was higher (Fig. S6). In the CON, females had higher protein content than the males in all generations except for the CON_july_, where males’ protein content was higher. The *horizontal* test in the ETP showed the *dev* effect was significant across the generations (*F* = 12.7, P < 0.001), with ETP_june_ and ETP_july_ having higher protein content than ETP_april_ or ETP_may_ (P < 0.05). The *horizontal* tests in the CON showed that the *dev* effect was significant (*F* = 10.6, P < 0.001) and showed a similar trend across generations as the ETP. The *vertical* tests showed there is no developmental effect for either the April or May generations. In the June and July generations, we see a significant *dev* by *sex* effect (*F* = 5.87, P = 0.027 and *F* = 5.58, P = 0.031, respectively), although neither generation show a significant difference in pairwise *post hoc* tests.

## 4. Discussion

As the environment changes in a predictable manner from spring (low thermal stress) to summer (high thermal stress) cactophilic *Drosophila* are predicted to modulate numerous traits, including behavioral, life history, and physiological traits (Marron et al., 2003; Behrman et al., 2015; Varpe, 2017). Using cues, many insects, can adjust their phenotype to successfully endure the summer or winter conditions (Masaki, 1980). Many cues (day length, temperature, humidity, food availability and crowding), often in combination, may serve as the catalyst to trigger changes in insect physiology, behavior, life history, etc (Nijhout 1999; Huestis and Lehmann, 2014). The key for these cues to act as such is that they are predictably tied to the oncoming harsh climate; however, they must be sensed prior to the climate becoming too stressful for the insect (Hodek, 2012). For example, such phenomena has been shown in *D. simulans* (Manenti et al., 2014), suggesting that unpredictably fluctuating environments are more stressful than predictably fluctuating or constant environments.

In multivoltine organisms, it may be the parental generation that senses the environment. The parental generation then affects the offspring phenotype, mediating changes that allow survival through both favorable and unfavorable periods. Allocation of energy and food resources into offspring, epigenetic signatures and choice of oviposition sites are all ways that parents may have direct effects on offspring success. Recent work on *D. mojavensis* has shown that transgenerational plasticity plays a significant role in larval heat resistance (Diaz et al., 2021). In the current study, transgenerational effects were not formally tested, however pursuing this avenue of research could prove to be promising.

While we hypothesized that given the environmental conditions experienced by *D. mojavensis* an aestivation phenotype would be expressed for the June and July treatments, a canonical aestivation response was not observed. Overall, fecundity of flies of ETP_june_ and ETP_july_ were as high as CON_june_ and CON_july_ when tested in the experimental temperature cycles. Although there were some differences in the number of eggs laid in the first five days as compared to the last five days (P < 0.001 for both), the fecundity of CON_may_ tested both at constant 25°C and at the experimental temperature was significantly depressed (Fig. 5). Given that the sharp decline in fecundity was only observed in the CON_may_ population, it is likely that this was due to an unaccounted environmental factor experienced by that generation only. Even if the CON_may_ data prevents a more definitive conclusion regarding a fecundity response from being drawn, the pattern of a lack of reduction in fecundity of the ETP populations in June and July strongly holds (Fig. 5).

While viability steadily increased over the generations of the CON tested in constant 25°C, it steadily decreased over the generations when tested in the experimental temperature cycles (Fig. S3). The ETP on the other hand, displayed the same pattern across generations in both temperature cycles, with the peak viability in ETP_may_. Longevity for ETP_may, june, july_ experiences a large drop when tested in the experimental temperature cycle but does not display this drop when tested at constant 25°C (Fig. 6). Additionally, although CON_may_ drops in number of days survived similarly to ETP_may_, CON_june, july_ live longer than ETP_june, july_ when tested in the experimental temperature cycles. Although this result could suggest that ETP_june, july_ flies are simply less robust in general than the CON_june, july_ flies, the metabolic pools for ETP_june_ do not suggest that, nor does the average longevity when tested in constant temperatures.

We did observe that, as predicted, that ETP_june_ and ETP_july_ flies had higher heat stress resistance, as did their offspring (Figs. 7, 8). Nutritional stress resistance was only measured in the July generation. It was expected that the ETP_july_ would be both more desiccation resistant and more nutritional stress resistant. However, this pattern was not observed; only desiccation resistance was higher in ETP_july_ (Figs. S4, S5). Given the importance of stores of lipids in starvation resistance (Marron et al., 2003), we can understand why the ETP_july_ flies were ill-adapted for nutritional stress, ETP_july_ flies had the lowest levels of all metabolic contents observed in this experiment. Additionally, the nearly ten hour lower average survival of ETP_july_ than ETP_june_ in constant 25°C can be explained in the same manner; for desiccation resistance, glycogen stores are extremely important (Marron et al., 2003), and ETP_june_ had higher glycogen stores than any other month.

Activity level peaks shifted over the four generations in the experimental temperature cycles in somewhat predictable manners, showing an advantageous response of both developing and being tested in the experimental temperature cycle. ETP_june_ and ETP_july_ flies had lower total diurnal activity and their peak activity was shifted earlier in the day, matching the coolest period of daylight (Fig. 4). The CON_june_ and CON_july_ flies tested in the experimental temperature cycle also shifted their peaks, however, they did not lower their overall activity, which would likely be detrimental to the fitness of *D. mojavensis*, given the high heat of the July temperature cycle. As we do not account for any potential behavioral compensation, we are not able to fully apply our activity results more generally to the natural population, but it does provide a framework for future work.

In this study we focused on altering both temperature and photoperiod across our generations, as this has been previously suggested to be the more ecologically relevant approach (Angilletta et al 2019). However, other environmental changes could serve as important factors, such as humidity and food availability. Additionally, these flies were maintained on an artificial food substrate, banana-molasses media, instead of the cactus necroses they would normally utilize, which could also contribute to the unexpected finding of a lack of reduction in reproduction in July. Cactus necroses are generally higher in protein content, lower in sugar content, and similar in triglyceride than the banana food used (L. Matzkin, unpublished).

In the case of an aestivation response, the expected trade-off would be between reproduction and longevity. In this study, we found that generations of the ETP closer to peak summer oviposited equal numbers of eggs overall but died earlier. *D. mojavensis* in the later generations of the ETP had increased stress resistance. Given our findings, the data reflect that the phenotypes are responding plastically to the environmental conditions, allowing *D. mojavensis* to persist in the Sonoran Desert summers. Given our experimental design, direct acclimation and transgenerational acclimation are confounded, and further research will need to be done in order to separate these effects.

Alternatively, our results may suggest that it is not the adults which are surviving the stressful summer, but instead an immature stage. Perhaps a trade-off between the stress resistance and longevity allows the flies to continue to lay eggs at the same rate, rather than go into an aestivation stage. If that were the case, it would mean a pre-adult stage of *D. mojavensis* must be able to survive the harsh Sonoran Desert summers. Recent findings that transgenerational effects are strongest on the larval stage (Diaz et al., 2021) suggest that the offspring of the summer generations of these flies are more acclimated to the summer heat. This expectation was confirmed in our finding that larva of later generations of the ETP were more heat resistant. In the Sonoran Desert, it is not only thermal stress, but also water stress that challenge individuals; however, precipitation is dramatically increased near the end of the summer during the monsoons. If an immature stage of *D. mojavensis* can survive until this time, they may have a better chance of survival. A possible model for *D. mojavensis* is that the summer conditions create a bottleneck in the population, as the survival rate is likely very low. The surviving flies may continue to oviposit and those pre-adult stages that can survive until precipitation arrives and temperatures cool go on to seeding the next generation.

## 5. Conclusion

In this study we tested for the occurrence of phenotypic changes in the cactophilic *D. mojavensis* between the spring and summer seasons in the Sonoran Desert. Although the flies did not go into a canonical aestivation stage as originally predicted, there were significant adjustments to the flies’ physiology, life history and behavior. These results leave us with some interesting hypotheses that will provide grounds for further experimentation. Specifically, we hypothesize that the necrotic host could provide essential nutrients which allows a subset of adults to survive the summer and/or that potentially non-adult stages are surviving the thermally stressful summer period. Current genomic and population genomic studies in this and related cactophilic *Drosophila* will further help illuminate the mechanism of adaptation to harsh desert conditions in these insects. Additionally, focusing on the regulation of mRNA might allow us to begin to infer the mechanistic causes for phenotypic changes observed.

## Abbreviations

CON: Control, population raised at constant 25°C. Subscript following denotes the generation.
ETP: Experimental Control Population, population raised at experimental temperature cycles. Subscript following denotes the generation.
*dev*: developmental temperature cycle in which the group was raised
*test*: temperature cycle in which the phenotype was measured.

## Credit authorship contribution statement

**Tierney M. Shaible:** Data curation, Formal analysis, Investigation, Methodology, Writing – original draft, Writing – review & editing. **Luciano M. Matzkin:** Investigation, Conceptualization, Funding acquisition, Methodology, Writing – review & editing.

## Acknowledgements

We would like to thank Carson Allan for his assistance in this project. This work was supported by the University of Arizona and an NSF grant (IOS-1557697) to LMM. We thank the Entomology and Insect Science Graduate program at the University of Arizona for support. Lastly, we like to thank the United States National Park Service at Organ Pipe National Monument for allowing the original collection of the *Drosophila* utilized to establish the stocks used in this study.

