## Supplemental figures for "Physiological and life history changes associated with seasonal adaptation in the cactophilic *Drosophila mojavensis*"

**Supplemental Material**

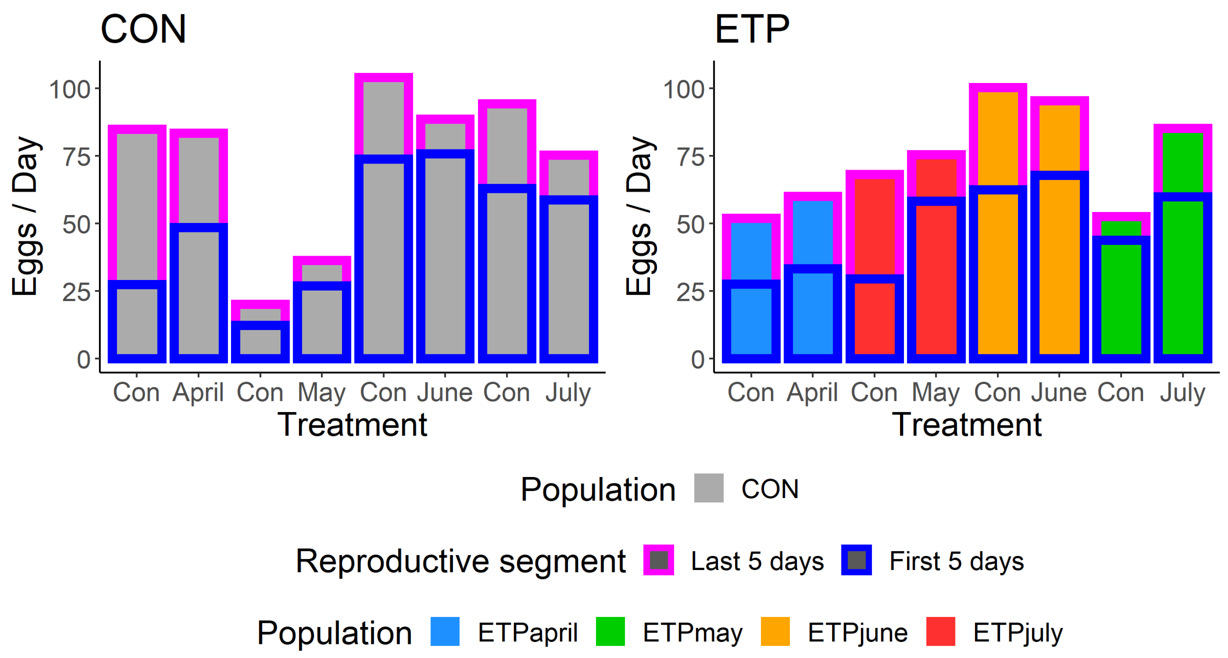


**Fig S1.** Mean fecundity measurements (eggs/day per female). The average number of eggs laid per day for the first five days is outlined in dark blue, while the average number of eggs laid per day for the last five days is outlined in pink. See tables S4 and S5 for additional information on subgroup differences.

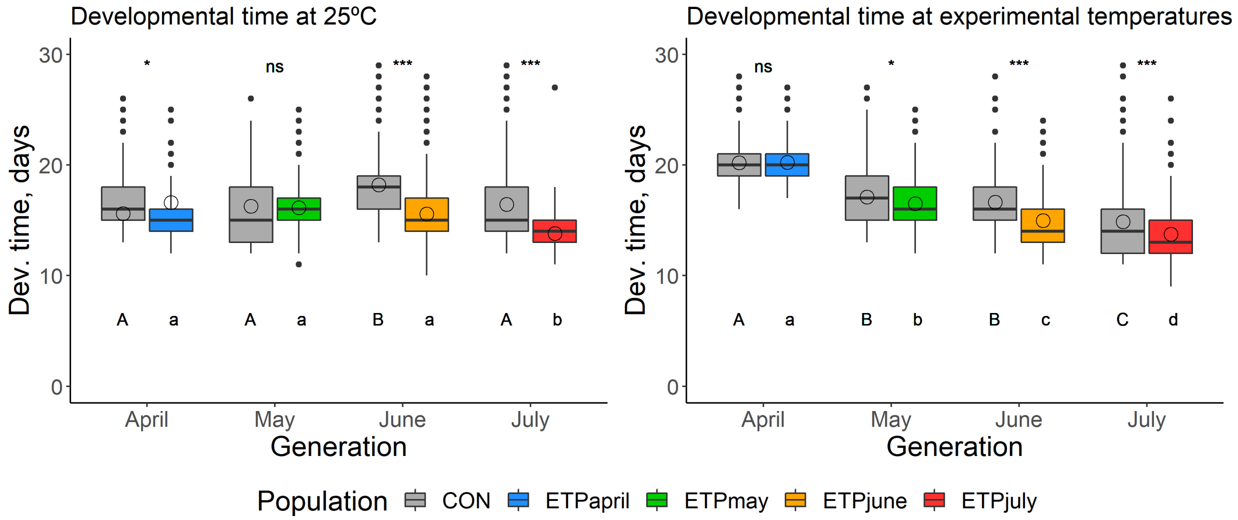


**Fig S2.** Developmental time measured as the number of days from oviposition to adult eclosion. Box plot horizontal lines indicate the first, second, and third quartiles and whiskers show the extreme upper and lower values within 1.5 times the interquartile range (IQR). Solid circles are values beyond 1.5× IQR. Open circles denote the average for the group. Asterisks indicates significance (*p < = 0.05, **p < = 0.01, ***p < = 0.001) between CON and ETP populations of the same generation. Uppercase letters indicate significant differences across generations of the CON population, while lowercase letters indicate significant differences across generations of the ETP.

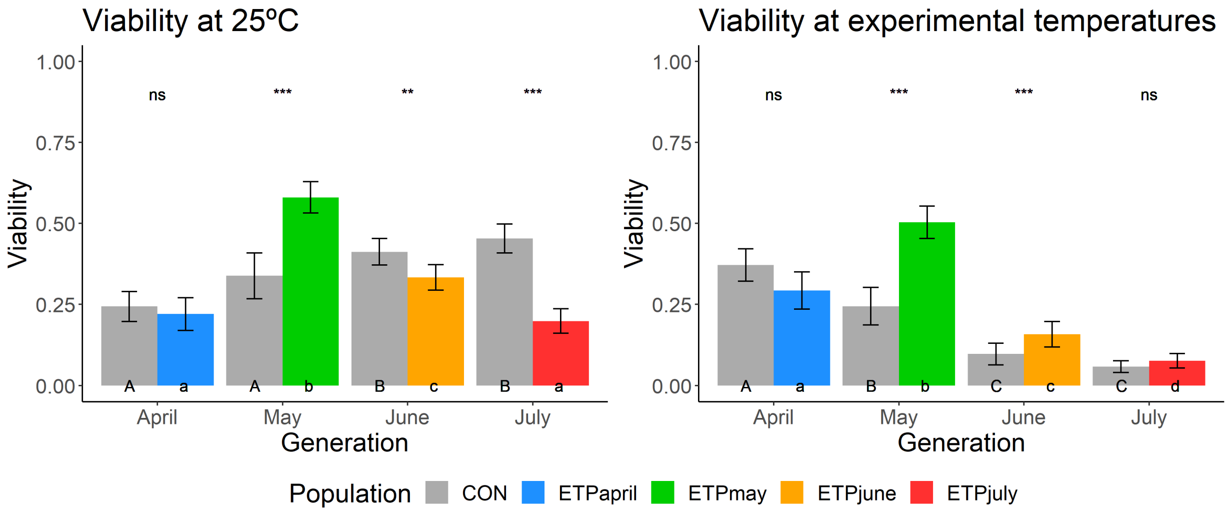


**Fig S3.** Mean egg-to-adult viability (± std. error) across the populations and treatments. Asterisks indicates significance (*p < = 0.05, **p < = 0.01, ***p < = 0.001) between CON and ETP populations of the same generation. Uppercase letters indicate significant differences across generations of the CON population, while lowercase letters indicate significant differences across generations of the ETP.

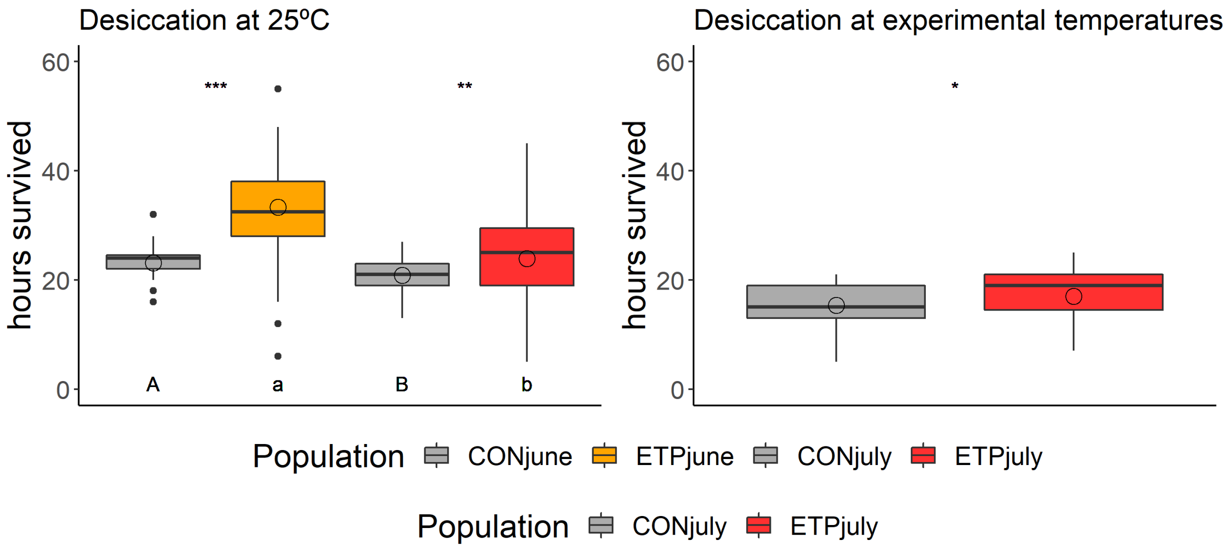


**Fig S4.** Desiccation (hours) resistance for the June and July generations. Box plot horizontal lines indicate the first, second, and third quartiles and whiskers show the extreme upper and lower values within 1.5 times the interquartile range (IQR). Solid circles are values beyond 1.5× IQR. Open circles denote the average for the group. Asterisks indicates significance (*p < = 0.05, **p < = 0.01, ***p < = 0.001) between CON and ETP populations of the same generation. Uppercase letters indicate significant differences across generations of the CON population, while lowercase letters indicate significant differences across generations of the ETP.

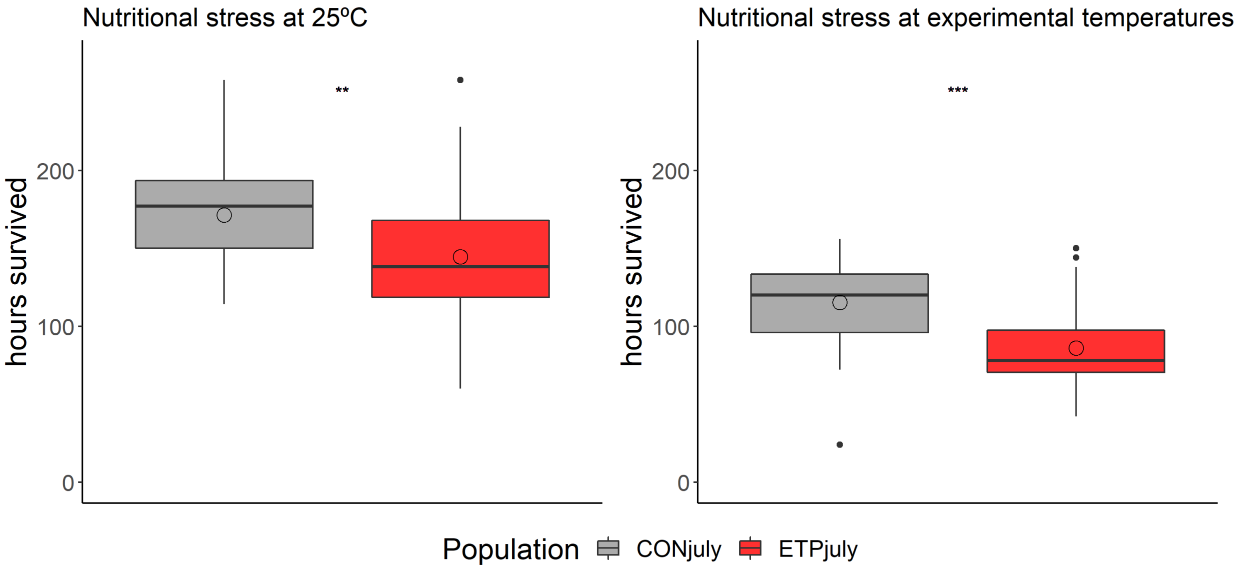


**Fig S5.** Nutritional stress survival (hours) for the July generation. Box plot horizontal lines indicate the first, second, and third quartiles and whiskers show the extreme upper and lower values within 1.5 times the interquartile range (IQR). Solid circles are values beyond 1.5× IQR. Open circles denote the average for the group. Asterisks indicates significance (*p < = 0.05, **p < = 0.01, ***p < = 0.001) between CON and ETP populations of the same generation.

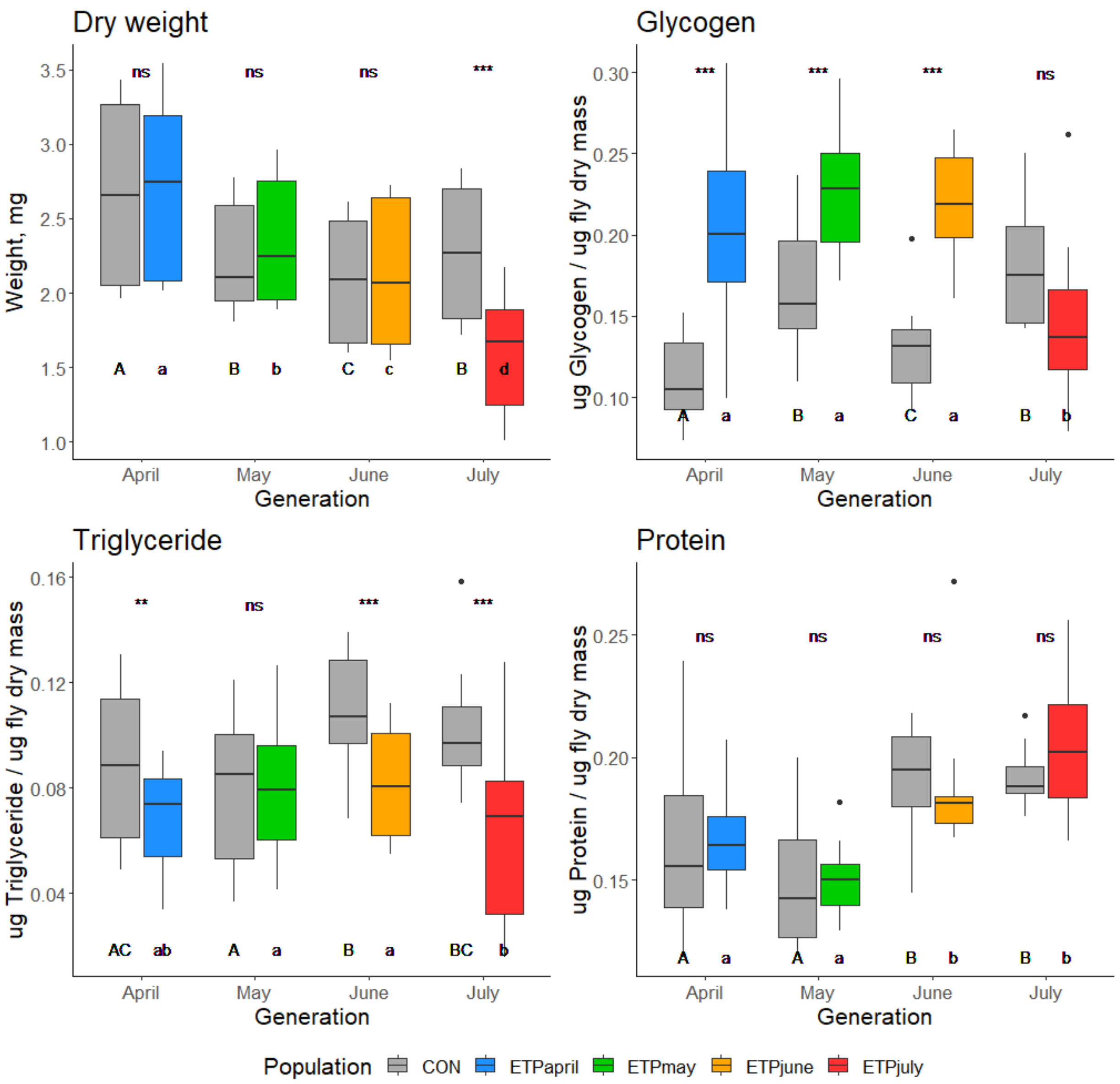


**Fig S6.** (A) Dry weight (mg) (B) Glycogen content (μg glycogen / μg fly dry mass) (C) Triglyceride content (μg triglycerides / μg fly dry mass) (D) Protein content (μg protein / μg fly dry mass). Box plot horizontal lines indicate the first, second, and third quartiles and whiskers show the extreme upper and lower values within 1.5 times the interquartile range (IQR). Solid circles are values beyond 1.5× IQR. Asterisks indicates significance (*p < = 0.05, **p < = 0.01, ***p < = 0.001) between CON and ETP populations of the same generation. Uppercase letters indicate significant differences across generations of the CON population, while lowercase letters indicate significant differences across generations of the ETP.
